# Capabilities of the CCi-Mobile Cochlear Implant Research Platform for Real-Time Sound Coding

**DOI:** 10.1101/2022.12.08.519687

**Authors:** Mahan Azadpour, Juliana Saba, John H.L. Hansen, Mario A. Svirsky

## Abstract

One important obstacle to optimizing fitting and sound coding for auditory implants is lack of flexible, powerful and portable platforms that can be used in real-world listening environments by implanted patients. The clinical processors and the typically available research tools either do not have sufficient computational power and flexibility or are not portable. In response to this need, the Center for Robust Speech Systems (CRSS) at the University of Texas at Dallas has developed CCI-Mobile, in collaboration with the Laboratory for Translational Auditory Research at New York University School of Medicine and the Binaural Hearing and Speech Laboratory at the University of Wisconsin-Madison. The CCI-Mobile platform provides unique flexibility to implement a variety of real-time sound coding algorithms in real-world environments, including algorithms that require synchronized binaural stimulation. In this paper, we will describe the overall architecture of the CCI-Mobile platform and provide practical considerations for designing real-time sound coding algorithms with this platform. CCI-Mobile is under development and future generations may provide further functionality, beyond what is described in this paper.

## 1. Introduction

Cochlear implant sound coding strategies have remarkably advanced over the past decades, paralleled by significant improvements in patient outcomes (Cullington and Zeng, 2008; Wouters *et al*., 2015). These advances owe to ongoing research on improving sound coding and signal processing algorithms that convert acoustic signal into electrical stimulation of the neural system. Valid evaluation of sound coding strategies and determining their ultimate potential frequently requires weeks or months of listening experience for cochlear implant users to adapt to the novel strategies. To be able to perform these evaluations, researchers need access to flexible, powerful and portable platforms that can be used in real-world listening environments by implanted patients. Cochlear implant researchers generally have access to portable clinical processors used by patients in daily life. However, the clinical processors and the fitting software have limited computational flexibility and power and the range of stimulation paradigms allowed by these devices is usually much restricted compared to the capabilities of the internal implant electronics. The research hardware/software tools provided by cochlear implant manufacturers are more flexible than the clinical processors and have played a critical role in advancing research in the auditory implant field (Litovsky *et al*., 2017). However, most of these research platforms cannot be used to implement and test real-time sound coding strategies and are not portable.

In response to this need, the Center for Robust Speech Systems (CRSS) at the University of Texas at Dallas has developed CCI-Mobile, in collaboration with the Laboratory for Translational Auditory Research at New York University School of Medicine and the Binaural Hearing and Speech Laboratory at the University of Wisconsin-Madison. CCI-Mobile is a portable/wearable real-time cochlear implant processing and stimulation platform for implant devices manufactured by Cochlear Corporation (Ghosh *et al*., 2022; Hansen *et al*., 2019). The CCI-mobile platform provides unique flexibility to implement a variety of real-time sound coding algorithms in real-world environments, including algorithms that require synchronized binaural stimulation. The goal of this paper is to introduce the features and capabilities of CCI-Mobile to the cochlear research community. We will describe the overall architecture of the platform and provide practical considerations for designing real-time sound coding algorithms with it. CCI-Mobile is under development and future generations may provide further functionality, beyond what is described in this paper.

## 2. CCI-Mobile platform

Figure 1 shows the overall diagram of CCI-Mobile platform and the communication between different components. In this paper, CCI-Mobile platform refers to the system of four components: Interface board, BTE (behind-the-ear) microphone, RF (radiofrequency) transmitter coils, and computing device. The core of CCI-Mobile is the interface board, which is connected with the other three components (BTE microphone and RF coils, computing device). Below we will describe the function of each component.

**Figure 1.**
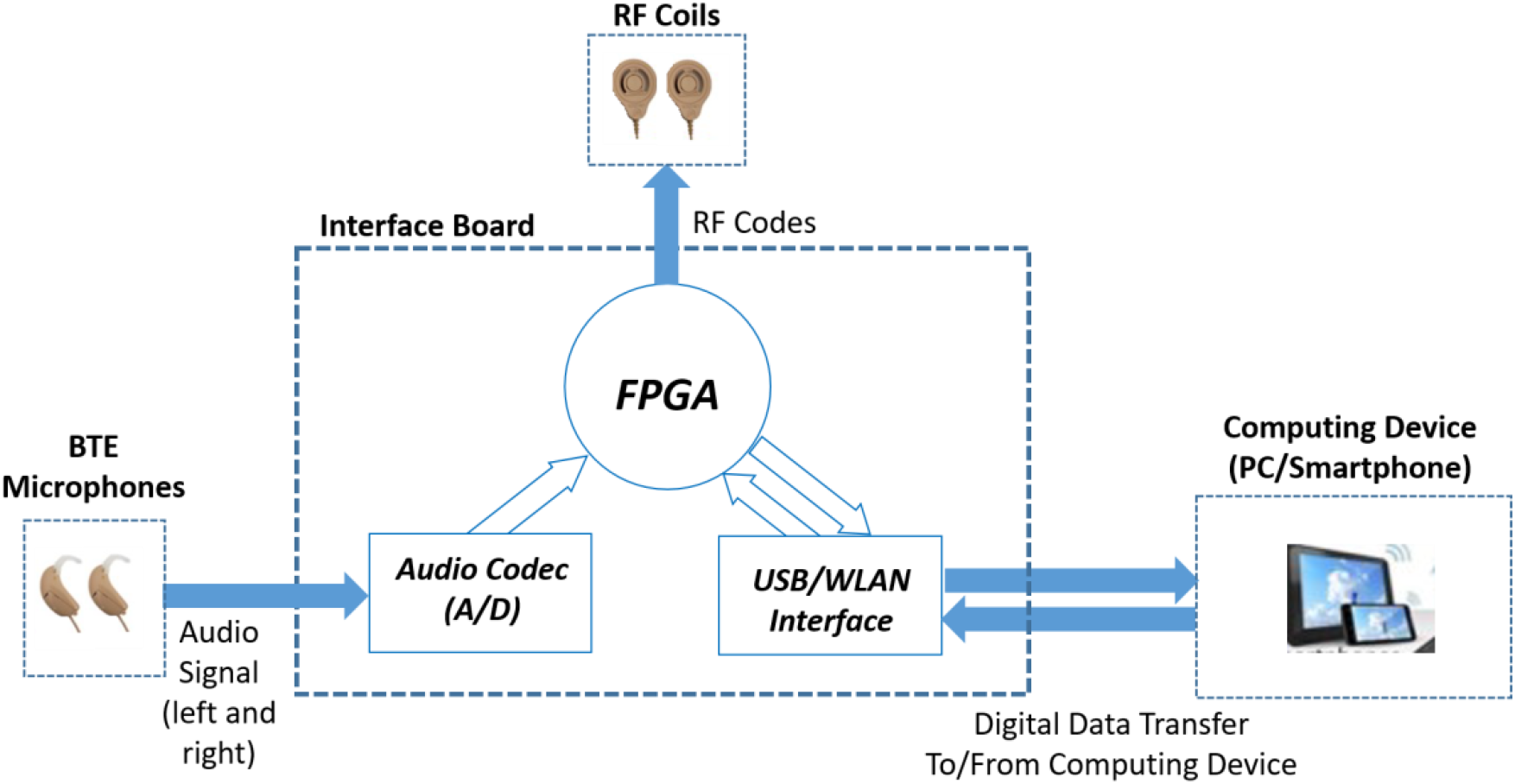
Overall diagram of CCI-Mobile platform.

### 2.1. Interface board

The CCI-Mobile interface board consists of three main modules: FPGA (field programmable gate array), audio codec, and USB/WLAN interface. The FPGA is the core of the interface board and manages communication with the external components (BTE microphones, RF coils and computing platform), as well as communication between the different modules within the interface board. Data transmission between the interface board and the computing platform is via USB and performed by the USB/WLAN interface module. Both wired USB and wireless WI-FI modes of data transmission are supported between CCI-Mobile and the computing platform (wireless connection is not currently fully functional). The audio codec module board acquires and digitizes synchronous audio from the left and right BTE microphones. Digital audio samples are quantized in two bytes at 16,000 samples/second, which is equivalent to the sampling rate of the commercial devices of Cochlear Corporation.

### 2.2. Computing device

Streaming with CCI-Mobile requires a Windows (PC or tablet) or Android (smartphone or tablet) computing device. The possibility to use an Android smartphone provides extreme portability compared to other available CI research platforms. The computing device communicates with the CCI-Mobile interface board via USB. Two types of data are transmitted between the interface board and the computing platform. One is digital audio samples, which are acquired by the audio codec and provided to the computing platform for signal processing (Figure 1). The other type of data is stimulation sequence data. Stimulation sequence data is generated by the computing platform and sent to the FPGA for RF generation and implant stimulation. The stimulation data can be determined by a sound coding strategy using the audio signal as input, or it can be determined by the user (e.g. for controlled direct stimulation). CCI Mobile comes with Matlab and Java software libraries to allow users to establish serial communication with the interface board and send/receive digital data. The Matlab libraries are for Windows PC/tablet devices. The Java libraries are for Android smartphone/tablet devices.

## 3. Streaming with CCI-Mobile

The CCI-Mobile streams stimulation to the patient’s implants via RF communication. The FPGA converts the stimulation data received by the computing device to RF and streams to the patient’s implants. The stimulation data is buffered into 516-byte packets for transmission to the FPGA. The data buffer contains stimulation parameters corresponding to an 8ms time frame for both left and right implants. The detail of stimulation data is described in the following section. Once streaming for 8ms is completed, the FPGA uses the updated buffer for stimulation in the following 8-ms period. To assure continuous streaming to the implant, the computing platform needs to update the FPGA stimulation data buffer every 8ms.

### 3.1. Stimulation data

The stimulation parameters for each 8-ms stimulation frame are determined by the computing device. Table 1 shows stimulation data for each 8-ms stimulation buffer. The computing device can use input audio form BTE microphones or stored audio (in a wave file for instance) to generate stimulation data for 8-ms time frames. Figure 2 shows data communication between the computing platform and the CCI-Mobile interface board, and the stimulation information for each 8-ms encoded in the 516-byte data buffer. Below we describe the details of stimulation data. Please note that the parameters described here refer to the current version of the CCI-Mobile.

**Table 1.** Pulse data parameters per 8-ms stimulation frame

| Parameter | #Bytes | Range |
| --- | --- | --- |
| Number of Pulses | 1 | 1 - 116 |
| Electrodes | 232 (116 for left and right implants) | 1 - 22 |
| Current Levels | 232 (116 per implant side) | 0 – 255 (CL) |
| Pulse Width | 4 (2 per implant side) | 8 – 400 (us) |
| Interphase Gap | 2 (1 per implant side) | Only 8 (us) |
| Stimulation Mode | 2 (1 per implant side) | Indicates internal chip<br>(CIC4 : 28, CIC3 : 30) |
| No-Stimulation Period | 4 bytes (2 per implant side) | 8 – 8000 (us) |

**Figure 2.**
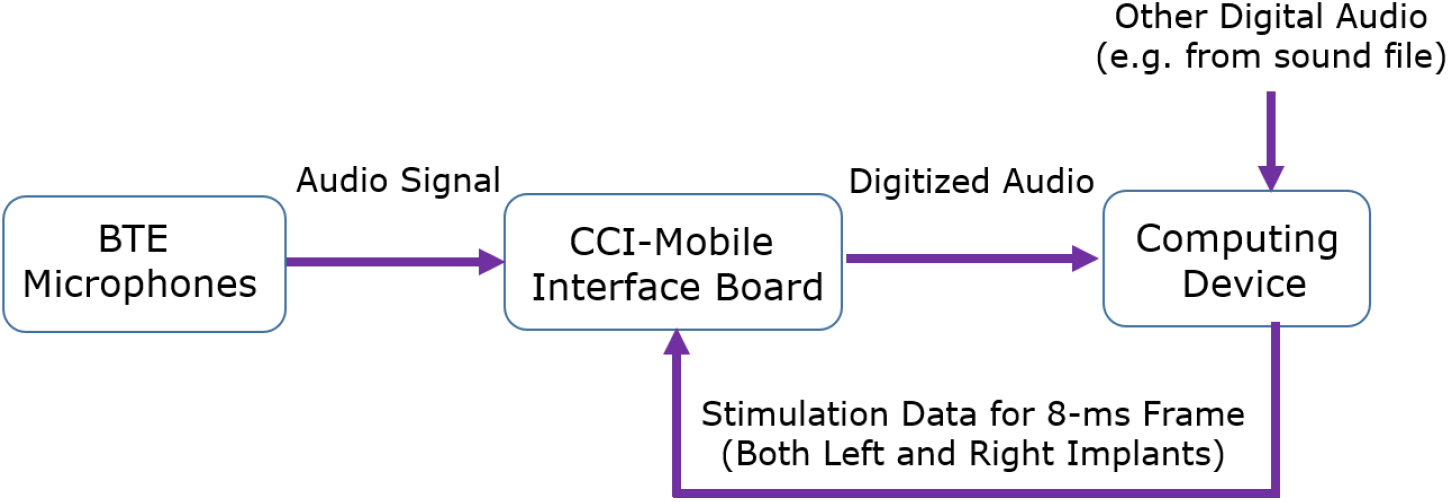
Audio and pulse data communication between CCI-Mobile microphones, interface board and the computing device.

#### Phase duration

The duration of biphasic pulses in each 8-ms frame is encoded by two bytes for each left and right implant. Phase duration must be the same for all the pulses within an 8-ms stimulation frame. In the current FPGA firmware version it is not possible for different electrodes to use different pulse widths within each 8-ms frame.

#### Interphase gap

the gap between the cathodic and anodic phases of biphasic pulses is fixed at 8 microseconds in the current FPGA firmware version and cannot be adjusted.

#### Stimulation mode

only the monopolar MP1+2 mode is supported in the current FPGA firmware version. Other modes including bipolar and common ground are not currently supported. The stimulation mode parameter is used to specify the type of the internal chip of the patient’s implant, i.e. CIC4 or CIC3.

#### Electrodes

Electrode numbers for pulses that occur in each 8-ms frame are specified by 116 bytes. Electrode for each pulse is specified by a byte. The maximum number of pulses in each 8-ms frame is 116 for each left and right implant, which corresponds to maximum 14,400 pulses/second that can be generated by the CCI-Mobile.

#### Current levels

The current levels for each of the pulses that occur in each 8-ms frame is specified by one byte. There are 116 bytes reserved for current levels in each 8-ms frame for each implant side.

#### Number of pulses

This parameter specifies the number of pulses that occur in an 8-ms stimulation frame.

#### No-stimulation period

The period of no-stimulation from the end of one pulse to the beginning of the next pulse is encoded by two bytes. The FPGA implements the no-stimulation period (in microseconds) at the end of each biphasic pulse.

### 3.2. Pulse timing

The no-stimulation period parameter determines the pulse-to-pulse period within a frame, which is related to pulse rate. The onset-to-onset timing between each pair of adjacent pulses within a frame is equal to the sum of “no-stimulation period” and the duration of biphasic pulse (i.e. 2 x “phase duration” + “inter-phase gap”). The current FPGA firmware starts pulses in each 8-ms stimulation frame from the beginning of the frame (time zero). With constant inter-pulse period across frames, the interval between pulses across consecutive frames will be equal to the inter-pulse period. This arrangement guarantees homogeneous pulse timings both within and between frames. However, since the no-stimulation period is fixed for the entire 8-ms frame, the users do not have control over the timing of individual pulses within a frame. This restricts the ability to set the timing of individual pulses, thus limiting the range of coding strategies that can be implemented with CCI-Mobile, e.g. fine-structure coding strategies that require precise timing control, or binaural strategies that encode interaural timing cues.

### 3.3. Bilateral Synchronization

CCI-Mobile is a unique real-time CI stimulation platform that allows synchronous stimulation between left and right implants. The 8-ms stimulation time frames are synchronized for the left and right implants, meaning that pulses begin at the same time for the two implant sides. This feature, however, restricts encoding of interaural time differences (ITD) between left and right implants. For precise ITD encoding, it might be desirable to apply ITDs for all pairs of pulses in the left and right implants. Nevertheless, it is still possible to implement interaural time differences by careful selection of stimulation parameters for each implant side, specifically the “no-stimulation period” and “number of pulses” parameters. ITD precision can in theory be improved by scaling the actual number of pulses per frame by an integer factor, and setting amplitude values only for the desired pulses. It is expected that future versions of the FPGA firmware will overcome the pulse timing limitations described above.

## 4. Real-time sound coding with CCI-Mobile

Sound coding is the process of converting audio signal into electric stimulation of the neural system. CCI-Mobile is capable of running sound coding strategies in real-time. This unique capability is crucial for providing long-term real-world exposure to novel sound coding strategies or evaluating strategies in real-world environments. Similar to standard clinical strategies, the sound coding algorithms with CCI-Mobile are performed on non-overlapping 8ms audio input frames. However, there are major implementation considerations due to the architecture of the FPGA stimulation frame and pulse data buffer. The CCI-Mobile sound coding strategies are required to determine pulse data for an entire 8ms stimulation frame, and update the FPGA data buffer every 8ms to assure continuous streaming to the patient’s implant. Figure 3 shows an overall diagram of real-time sound coding with CCI-Mobile.

**Figure 3.**
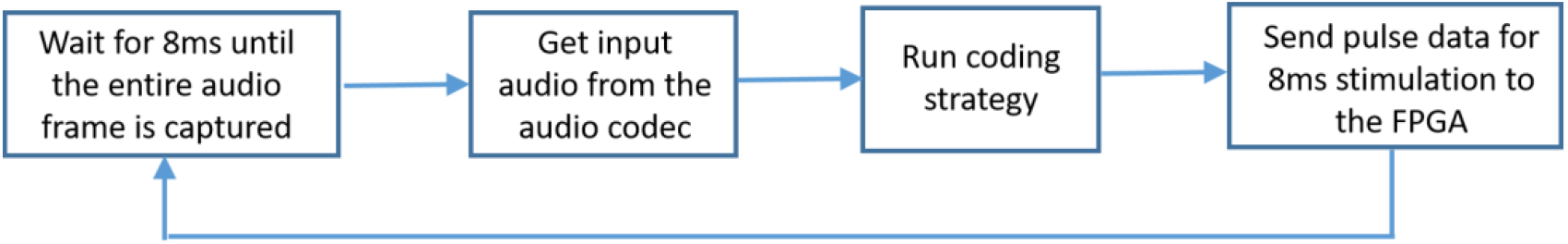
Stages for real-time sound coding with CCI-Mobile, from audio capturing to stimulation

### 4.1. Real-time ACE strategy with CCI-Mobile

CCI-Mobile users have access to open-source implementation of ACE (Advanced Combination Encoder) strategy, which is the default clinical sound coding strategy of Cochlear Corporation (Patrick *et al*., 2006). ACE selects only a subset of electrodes for activation in each stimulation cycle (Seligman and McDermott, 1995). The electrodes and the corresponding current levels are determined by an FFT-based envelope extraction approach applied to 8-ms blocks of audio signal (128 samples). Frequency channels are created from FFT and channels with the highest envelope energies are used for determining electric stimulation parameters. An overview of the major ACE processing stages are shown in Figure 4. The ACE output from one audio blocks determine stimulation for a cycle.

**Figure 4.**
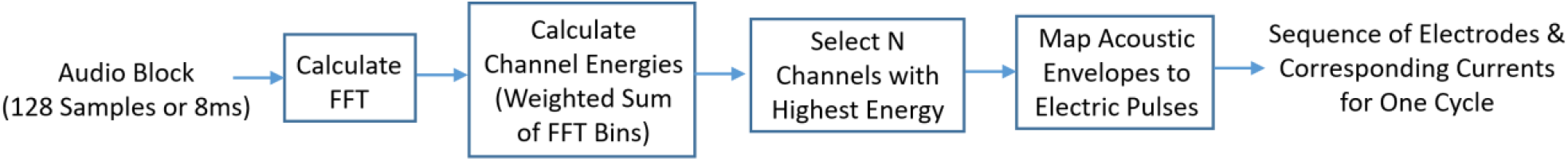
ACE processing strategy

Audio blocks are constructed from the input audio signal, by sliding a window of 128 samples (8ms) over the input audio frames. Figure 5 shows the process of constructing audio blocks from the input audio frames. The number of audio samples between consecutive blocks (block-shift) determines stimulation rate (cycles/second). The time between blocks (i.e. block-shift/audio sample rate) determines the period between stimulation cycles, which is the inverse of stimulation rate. For instance a block shift of 16 samples at 16,000 samples/second corresponds to 1ms cycle period or 1000 cycles/second.

**Figure 5.**
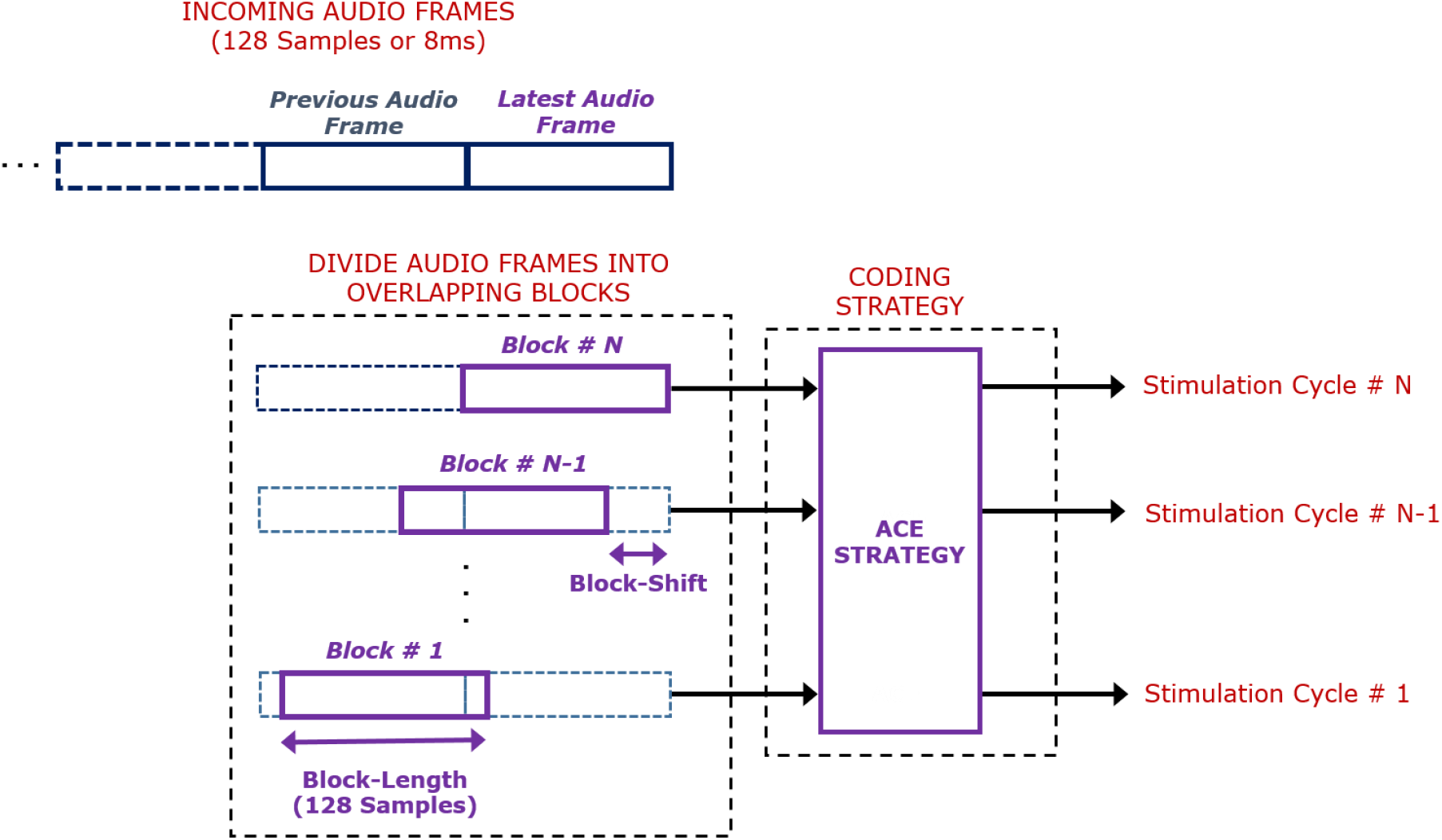
Construction of audio blocks from the input audio frames

The CCI-Mobile implementation of the ACE strategy mirrors the processing stages in the Nucleus Matlab Toolbox (NMT) provided by Cochlear Corporation. NMT provides a reasonably close simulation of the state-of-the-art clinical processing strategies. The CCI-Mobile software routines are in Matlab and Java, to use with PC and Android device respectively. CCI-Mobile software is open-source, available to the research community and can be readily used by the researchers or be customized for novel strategy designs. There is no intellectual property for the software routines.

### 4.2. Stimulation rate with CCI-Mobile

#### Direct implementation of pulse rate by pulse data parameters

The current FPGA firmware generates stimulation for an entire 8-ms stimulation frame such that the first pulse in each frame occurs at time zero from the beginning of the frame. This arrangement can result in non-uniform pulse presentation depending on the pulse rate (R), defined as the number of pulses per second. The number of pulses (N) per 8-ms frame is:

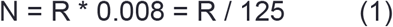

Uniform pulses across 8-ms frames would be achievable with integer values of N and by setting the time between pulses at 8/N milliseconds. An example for N=3 is shown in figure 6a. An integer N corresponds to stimulation rate R being multiple of 125 pulses/second. However, when N is not an integer (i.e. R is not a multiple of 125), the number of pulses per frame would need to vary across stimulation frames and will not be constant. Figure 6b shows pulses for R=400 pulses/second (corresponding to N=3.2). The top panel is the ideal arrangement of stimulation pulses across two consecutive frames. The bottom panel shows the actual pulses generated by the current CCI-Mobile firmware. The CCI-Mobile FPGA firmware can produce different number of pulses across frames, but does not currently allow changing the onset time of the first pulse in each frame. Future generations of CCI-Mobile may allow a wider range of stimulation rates by more flexibility in the timing of individual pulses. Currently, only multiples of 125 pulses/second can be accurately implemented with CCI-Mobile.

**Figure 6.**
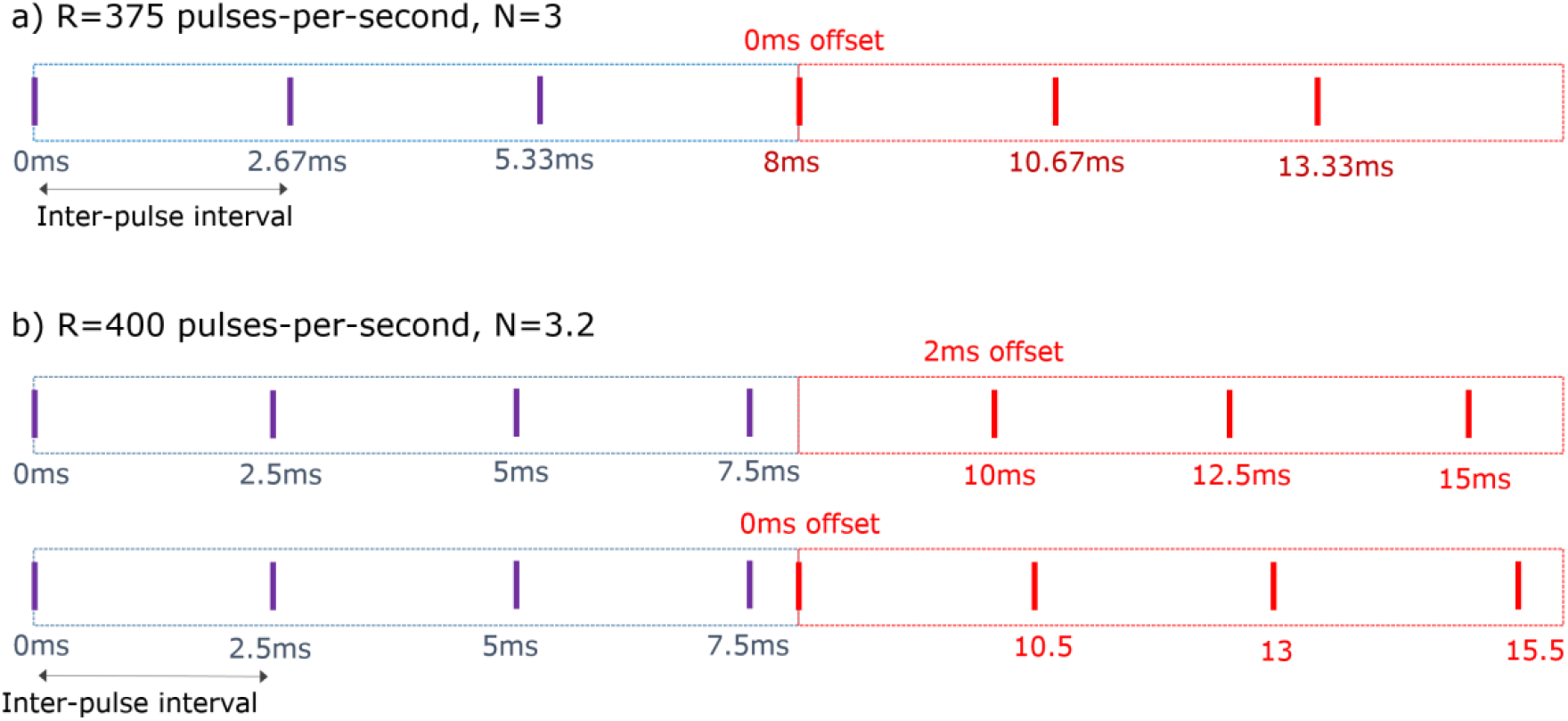
Schematics of pulse outputs for two different stimulation rates

#### Implementing stimulation rate in the sound coding strategies

The block-shift parameter of sound coding strategies determines the period between stimulation cycles, which is inverse of stimulation rate (Figure 5). Using equation (1), the number of cycles per frame (N) can be written as:

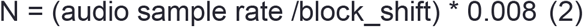

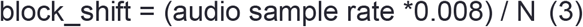

With 16,000 sampling rate:

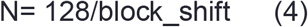

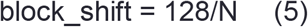

As discussed above and shown in Figure 6, uniform pulse distribution requires that N be an integer. To achieve integer N, block-shift must be a power of two. This will result in N itself being power of two as well. With this criteria, the only stimulation rates that would correctly result in uniform pulses are multiple and submultiples of 1000 cycles/second (i.e. 125, 250, 500, 1000, 2000, etc.). This stimulation rate criterion for ACE strategy (power of 2 multiples of 125) is more restricted than the rate criteria for direct stimulus generation, where any multiple of 125 would be acceptable.

We propose a work around to overcome limitations for implementing stimulation rate with sounding strategies of CCI-Mobile. The workaround is by resampling the audio signal and constructing audio blocks from resamples audio. The resampling procedure below will allow a wider range of integer cycles/frame (N) that can achieved with implementable integer block-shift parameter.

We propose the following steps for constructing the sliding audio blocks in the sound coding strategy:

1. Up-sample the audio frames by the desired cycles/frame (N). Simply repeat each sample in the audio signal N-1 times. This will be equivalent to scaling the audio sampling rate by N.
2. Set block-shift at 128. With the scaled audio sampling rate, the block-shift of 128 will maintain the desired cycles per frame at N (equation 2).
3. Scale the block-length by N.
4. Construct shifted audio blocks using the up-sampled audio frames and the new block length and block shift values.
5. Down-sample the constructed blocks by N. The down-sampling can be done by simply removing the repetitions that were inserted in the up-sampling stage. The length of the audio blocks will be scaled back to 128
6. Apply ACE strategy to the constructed audio blocks.

By fixing the block-shift parameter at 128, the proposed resampling technique will allow implementing stimulation rates that are any multiple of 125 cycles/second. This subset of stimulation rates is a larger subset that could otherwise be implementable with CCI-Mobile (i.e. only power of two multiples of 125 cycles/second such as 250, 500, 1000, 2000, and so on).

## 5. Pre-processing

The CCI-Mobile platform provides the capability to implement computationally intensive front-end processing algorithms that cannot be handled by the clinical processors. The ACE processing software provided with CCI-Mobile does not implement any front-end processing and it’s up to the users whether to implement front-end algorithms. The CCI-Mobile users would need to implement pre-emphasis when processing audio signals that are imported from a file.

The pre-emphasis is part of the ACE processing strategy to increase the likelihood of selecting high-frequency channels by emphasizing high-frequencies in the signal. In the toolbox provided by Cochlear, pre-emphasis is approximated with a high-pass one-order Chebyshev filter with cutoff at 1000Hz (Shekar and Hansen, 2021). Pre-emphasis filter is not required when the audio signal is recorded by the CCI-Mobile microphone. The frequency response of the CCI-Mobile microphone acts as a high-pass filter by attenuating lower frequencies in the audio signal.

## 6. Safety considerations

When utilizing the CCI-Mobile software code, the investigators are fully responsible for assuring patient safety. If required by investigator’s institution, ethics approval needs to be sought for studies conducted with CCI-Mobile. The CCI-Mobile FPGA enforces some of the important biological safety considerations for electrical stimulation. One important consideration is that it only allows symmetric biphasic stimulation such that the amount of charge during one phase is reversed during the other phase to prevent ionic charge imbalance and formation of electrochemical products at the electrode-tissue interface. This safety consideration is similar to what is done in clinical Cochlear™ processors. The internal Cochlear™ stimulator further enforces charge balance in the tissue by keeping all electrodes shorted after each electric pulse is generated (Litovsky *et al*., 2017).

As a platform developed for research purposes, it is mainly up to the user to make sure that stimulation is within the comfortable loudness limit of each individual implant patient. Stimulation level is normally adjusted to find a listener-specific range of comfortable levels. It is highly recommended that CCI-Mobile users implement software checks to assure that stimulation never exceeds levels that could be harmful to the patients. One example is preventing biological neural tissue damage.

The CCI-Mobile software codes have implemented safety guidelines to prevent neural damage (Shannon, 1992). These safety guidelines were based on animal neural damage data that were obtained after long period of electrical stimulation with electrodes that were in contact with the neural tissue (McCreery *et al*., 1990). Estimation of electric current that results in neural damage is given by:

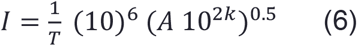

where *I* is electrode current in microamps, *A* is the electrode surface area in square millimeters,

*T* is pulse phase duration in microseconds. Equation (6) can be rewritten in term of electrode diameter D where 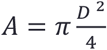

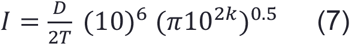

Constant *k* determines the conservativeness of the current estimate. Figure 7 shows the safety limits that are implemented in the CCI-Mobile software. The maximum safe current levels as a function of phase duration in microamps, as well as clinical units for CIC3 and CIC4 internal implant chips are shown in that figure. These maximum currents were calculated for a small electrode diameter D=0.3 mm corresponding to surface area A=0.07 mm^2^ and for *k* = 1.5, which result in a conservative estimate for clinical applications (Shannon, 1992).

**Figure 7.**
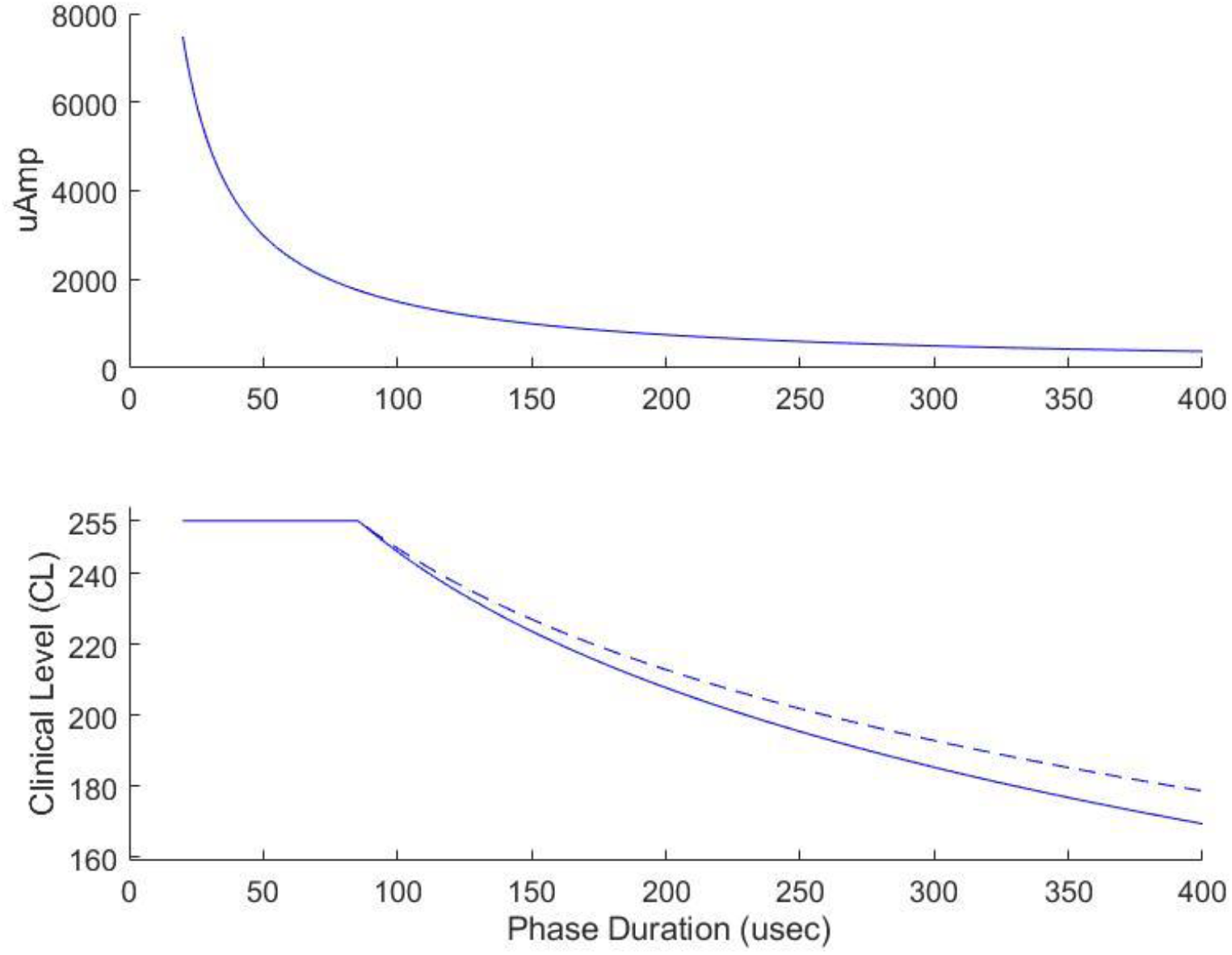
Safe current level limits that are implemented in the CCI-Mobile software codes. Shannon limits are shown for 0.3 mm electrode diameter and k=1.5.

## 7. Summary and future directions

In summary, CCi-MOBILE significantly extends the functionality available to the cochlear implant research community by allowing the development of novel stimulation strategies that go well beyond the capabilities of clinically available speech processors. The platform is open-source and portable, allowing CI users to experience a new stimulation strategy in the real world for as long as necessary to achieve asymptotic performance with the strategy. This process is known to require weeks, and thus it would be very difficult (if not impossible) to achieve with a non-portable, laboratory-based platform. Here we described the platform’s current functionality, as well as some if its limitations and possible workarounds.

It should be kept in mind that CCi-MOBILE is not a static finished product. For example, there are already plans to add ecological momentary assessment (EMA) functionality (Shiffman *et al*., 2008), and to allow for associated sample audio recordings from the environment obtained at the same time that EMA questionnaires are answered by the user. There are also plans to implement location-based (smart-space) connectivity with CCi-MOBILE. This would allow a smart room to broadcast information about its acoustic environment to CCi-MOBILE, which could then be used to modify speech processing on the fly. These examples speak to the fact that CCi-MOBILE is a platform that evolves based on user feedback and the needs and interests of the research community

